# Visual cortex cTBS increases mixed percept duration while a-tDCS has no effect on binocular rivalry

**DOI:** 10.1101/2020.09.07.285999

**Authors:** Dania Abuleil, Daphne McCulloch, Benjamin Thompson

## Abstract

Neuromodulation of the primary visual cortex using anodal transcranial direct current stimulation (a-tDCS) can alter visual perception and enhance neuroplasticity. However, the mechanisms that underpin these effects are currently unknown. When applied to the motor cortex, a-tDCS reduces the concentration of the inhibitory neurotransmitter gamma aminobutyric acid (GABA), an effect that has been linked to increased neuroplasticity. The aim of this study was to assess whether a-tDCS also reduces GABA-mediated inhibition when applied to the human visual cortex. Changes in visual cortex inhibition were measured using the mixed percept duration in binocular rivalry. Binocular rivalry mixed percept duration has recently been advocated as a direct and sensitive measure of visual cortex inhibition whereby GABA agonists decrease mixed percept durations and agonists of the excitatory neurotransmitter acetylcholine increase them. Our hypothesis was that visual cortex a-tDCS would increase mixed percept duration by reducing GABA-mediated inhibition and increasing cortical excitation. In addition, we measured the effect of continuous theta-burst transcranial magnetic stimulation (cTBS) of the visual cortex on binocular rivalry dynamics. When applied to the motor or visual cortex, cTBS increases GABA concentration and we therefore hypothesized that visual cortex cTBS would decrease the mixed percept duration. Binocular rivalry dynamics were recorded before and after active and sham a-tDCS (N=15) or cTBS (N=15). Contrary to our hypotheses, a-tDCS had no effect, whereas cTBS significantly increased mixed percepts during rivalry. These results suggest that the neurochemical mechanisms of a-tDCS may differ between the motor and visual cortices.

## Introduction

Anodal transcranial direct current stimulation (a-tDCS) is a non-invasive electrical brain stimulation technique that can modulate neural excitability and promote neuroplasticity. When applied to the visual cortex, a-tDCS can increase contrast sensitivity (1–4), improve visual acuity (5,6), and enhance perceptual learning (7,8) in patients with amblyopia, a neurodevelopmental disorder that affects binocular vision, as well as in controls. In addition to perceptual changes, reduced phosphene thresholds (7,9,10) and increased VEP amplitudes (4,7,11) have been reported following a-tDCS. Therefore, a-tDCS can induce physiological and neurochemical changes in the visual cortex that result in increased cortical excitability.

Although the specific mechanisms that underlie the effects of visual cortex a-tDCS are unknown, the effects of motor cortex a-tDCS are attributed in part to a reduction in cortical inhibition mediated by the neurotransmitter gamma amino-butyric acid (GABA). Specifically, magnetic resonance spectroscopy measures indicate that a-tDCS reduces motor cortex GABA concentration (12–18). We anticipated that a similar reduction in GABA concentration occurs when a-tDCS is applied to the visual cortex on the basis of previous results that are consistent with such an effect. For example, magnetoencephalography measurements made after visual cortex a-tDCS indicated an increase in occipital gamma activity that has been linked with reduced GABA-mediated inhibition (Wilson et al., 2018, but see Hanley et al., 2016; Marshall et al., 2016). In addition, visual phenomena that have been associated with neural inhibition such as an attenuated cortical response to inputs from the amblyopic eye in adults with amblyopia (3,4) surround suppression (22) and lateral inhibition (23) can be reduced by a-tDCS.

In contrast to a-tDCS, continuous theta-burst stimulation (cTBS), a form of transcranial magnetic stimulation that can also alter visual perception (24–27), has been found to increase GABA concentration in both the motor cortex (28) and the visual cortex (25). cTBS, therefore, would be expected to have the opposite effect to a-tDCS on percepts that are directly influenced by GABA-mediated inhibition.

Binocular rivalry dynamics have recently been advocated as a sensitive measure of GABA-mediated inhibition withing the human visual cortex (29). Binocular rivalry is a form of bistable perception wherein the brain alternately suppresses one eye over the other stochastically when each eye views a different image (30–33). Previous studies have found that binocular rivalry dynamics in young adults are correlated with visual cortex GABA concentration (34–36). Specifically, young adults with slower binocular rivalry alternation rates had higher primary visual cortex GABA concentrations (34,36). In addition, higher GABA concentrations are correlated with longer periods of perceptual dominance, defined as the period of time when either eye dominates perception during rivalry as opposed to mixed percepts when both eyes contribute to perception (36).

A causal relationship between GABA-mediated inhibition and binocular rivalry dynamics has also been reported. Single doses of clobazam (a GABA_a_ receptor agonist) or arbaclofen (a GABA_b_ receptor agonist) significantly increased perceptual dominance and reduced mixed percept duration during binocular rivalry compared to a placebo (29). Additionally, reduced inhibition and increased excitation induced by the acetylcholine agonist donepezil was recently found to reduce perceptual dominance and increase mixed percept duration during binocular rivalry (37). Given this evidence, we used binocular rivalry mixed percept duration as a measure of cortical inhibition.

Whether a-tDCS reduces visual cortex GABA concentration as it does in the motor cortex is not yet known. Our study aimed to address this question. We hypothesized that visual cortex a-tDCS would reduce visual cortex GABA concentration resulting in increased mixed percept durations during binocular rivalry. We further hypothesized that visual cortex cTBS, that has been found to increase visual cortex GABA concentration (25), would have the opposite effect.

## Materials and Methods

### Participants

A total of thirty young adults with normal or corrected-to-normal vision (0.1 LogMAR or better in each eye) participated in one of two within-subject, sham-controlled experiments: an a-tDCS experiment (n = 15, mean participant age 25, median age 24, range 22-30, 11 female) and a cTBS experiment (n = 15, mean participant age 24, median age 24, range 22-29, 7 female). Participants with abnormal binocular vision and those taking psychoactive drugs were excluded. All participants were informed of the nature of the study before participation and provided written informed consent. The project was approved by the University of Waterloo Research Ethics Committee (ORE #30537).

### Visual Stimuli

Dichoptic, orthogonally oriented (45° and 135°) sinusoidally modulated red/green gratings (0.5 cycles per degree, 6.1 degrees of visual angle) were presented on a 24-inch Asus 3D monitor. Participants wore frame sequential shutter glasses to view the stimuli. The contrast of the gratings was matched using a Chroma Meter CS-100^®^ photometer (mean luminance: red = 8.4 cd/m^2^; green = 32.9 cd/m^2^). Stimuli were viewed from 57cm using a chin rest. Participants reported perceiving the 45° grating only, the 135° grating only or a mixture of both (piecemeal or superimposition percepts) by holding down a computer keyboard key and switching keys as the percept changed.

### Anodal Transcranial Direct Current Stimulation

Two 5×7 cm electrode sponges were placed on the scalp, the anode at international 10-20 electrode system position Oz and the cathode at Cz. Each tDCS electrode was placed inside a saline-soaked sponge. A-tDCS was delivered at 2mA for 15 minutes in addition to a 30-second ramp-up and 30-second ramp-down period using a NeuroConn DC-Stimulator MC-8. The sham condition consisted only of the ramp-up and ramp-down periods. Participants were masked to the stimulation condition. The experimenter could not be masked due to resource limitations; however, session order (active first or sham first) was randomly sequenced prior to the start of data collection. For both active and sham conditions, six 60-second trials of binocular rivalry were completed before, during, 5 minutes, and 30 minutes post stimulation (Fig 1A).

**Fig 1:**
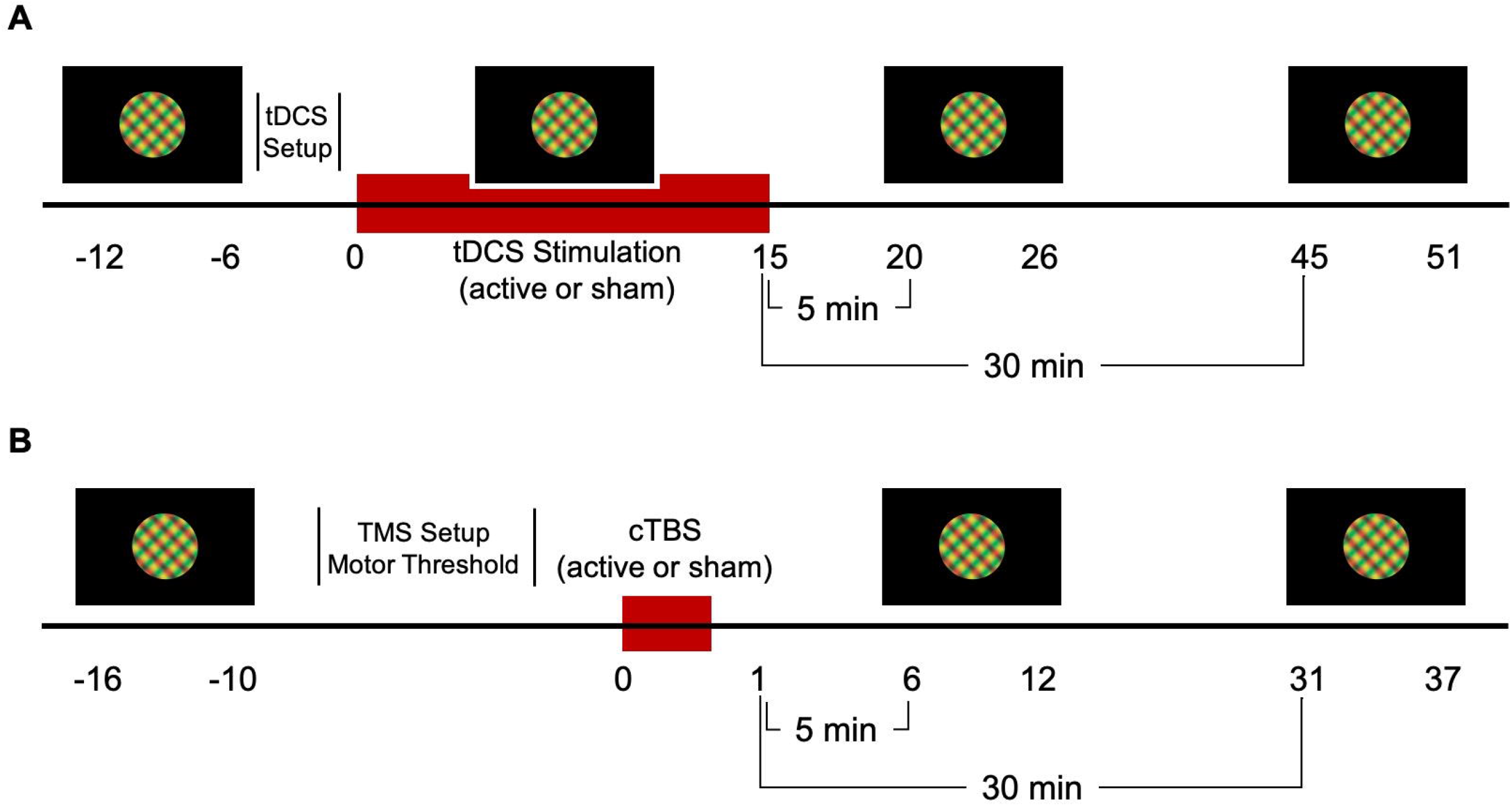
Protocol of the a-tDCS (A) and cTBS (B) experiments. Binocular rivalry dynamics were recorded for 6 minutes at baseline (pre), during, 5 minutes post and 30 minutes post a-tDCS. A-tDCS electrodes were placed on the head following the baseline measure. Similarly, for cTBS, binocular rivalry dynamics were recorded pre, 5 minutes post and 30 minutes post stimulation. Motor thresholding was completed on the first visit following the baseline measure.

### Continuous Theta Burst Stimulation

Stimulation was delivered using a MagVenture MagPro X100 stimulator (MagVenture Farum, Denmark) with BrainSight frameless neuro-navigation software (Rogue Research Inc., Montreal, Canada). Active motor thresholds (AMTs) were used to calibrate visual cortex cTBS intensity. The procedure for determining AMT involved placing a surface electrode on the belly of the first dorsal interosseous muscle tendon (left or right based on hand dominance) and a second electrode on the lateral bone of the wrist. The electromyographic (EMG) response was monitored using Brainsight software as the participant was asked to steadily press their pointer finger against the arm of their chair to generate a motor evoked potential (MEP) of 100µV. A single pulse of TMS was systematically delivered to different points of a contralateral motor cortex stimulation grid (3 by 3 cm) beginning at 40% of the maximum stimulator output (MSO) until the region hotspot—defined as the stimulation location corresponding to the maximum TMS-induced MEP amplitude—was located (38,39). Using the Rossini-Rothwell algorithm for determining AMT, single pulses were then delivered to this region while increasing the intensity by 1% until a peak-to-peak amplitude of 200µV was generated for 5 out of 10 pulses (50%) (40).

For visual cortex cTBS, the coil was placed over the occipital pole, identified as 2 cm above the inion, 0 cm lateral. Stimulation involved 600, 20 ms pulses delivered in 50Hz bursts for 40 seconds at 100% of the participant’s AMT. The control condition used the same protocol with a sham coil. Both the participant and experimenter were masked to the stimulation condition (active and sham condition codes were given to the experimenter by another researcher). Binocular rivalry dynamics were recorded for six 60-second trials before, 5 minutes post, and 30 minutes post stimulation (Fig 1B).

### Analysis

The duration of mixed perception during binocular rivalry was calculated in seconds per 60 second trial. We also analysed binocular rivalry ocular dominance index ((time viewing dominant eye percept – time viewing nondominant eye percept)/total time excluding mixed percepts) and alternation rates (any change in perception). Measures were averaged across all six trials separately for each participant. The dominant eye was defined as the eye with the longest pre-stimulation viewing time at the initial visit.

A repeated measures ANOVA with factors of Condition (active vs. sham) and Time (a-tDCS: pre vs. during vs. 5min post vs. 30min post; cTBS: pre vs. 5min post vs. 30min post) was conducted separately for mixed percept duration, ocular dominance index and alternation rate for each stimulation type. Post-hoc testing of significant interactions was conducted using t-tests.

For one tDCS participant, the 5 minutes post stimulation data for the sham condition was irretrievably lost. For one TMS participant, baseline data and 5 minutes post stimulation data for the sham condition were irretrievably lost. The missing data points were imputed using the mean value of the other 14 participants (41).

## Results

### Anodal Transcranial Direct Current Stimulation

No significant effects of a-tDCS were observed for any measure of binocular rivalry dynamics (p > 0.05). Fig 2 illustrates mixed percept duration, ocular dominance index and alternation rate for the active a-tDCS and sham conditions.

**Fig 2:**
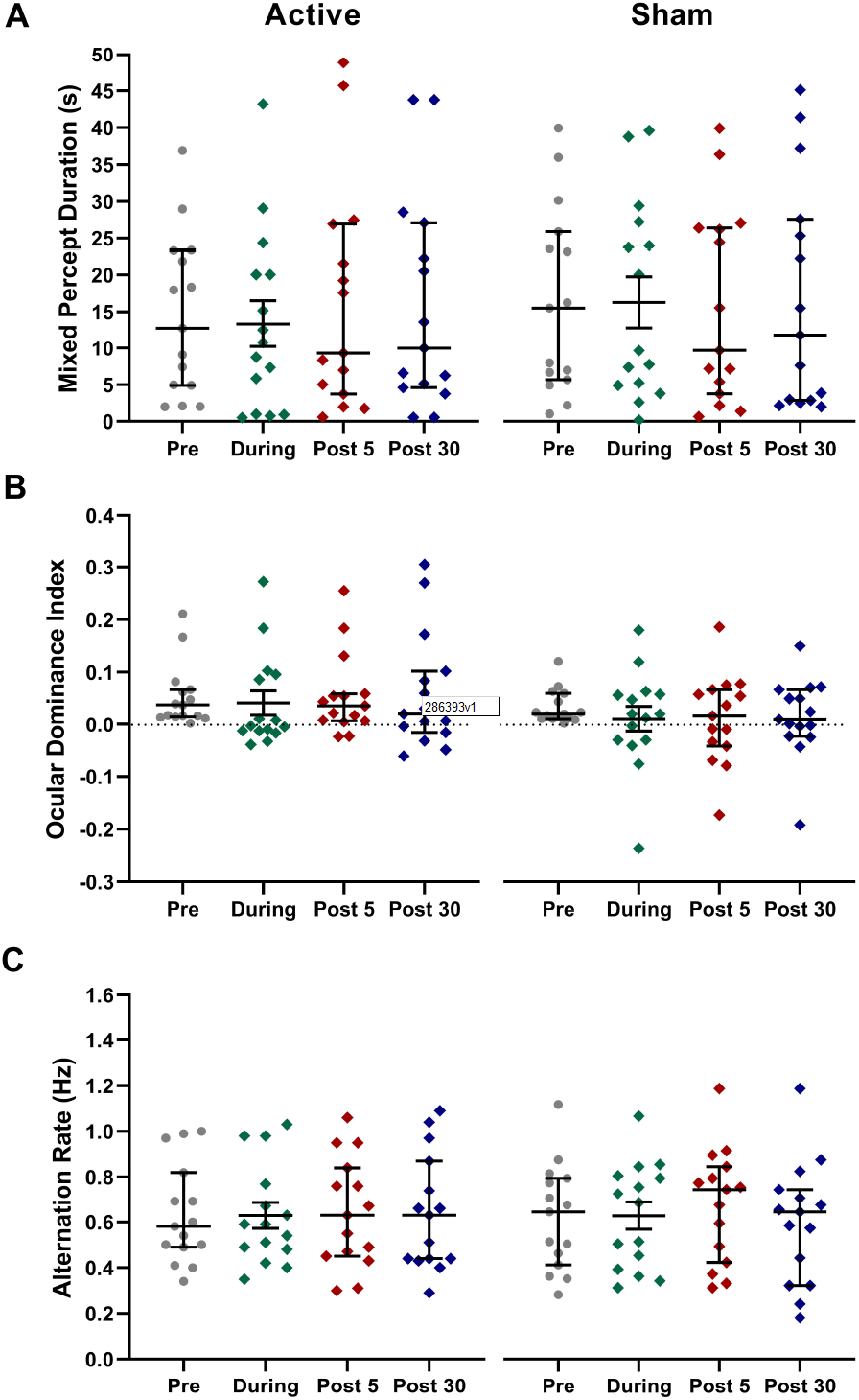
Average time spent in mixed percept (A), ocular dominance indices (B), and alternation rates (C) for 15 participants pre, during, 5 minutes and 30 minutes post a-tDCS. Error bars = SEM. No statistically significant effects were observed.

### Continuous Theta Burst Stimulation

cTBS significantly increased the duration of mixed percepts relative to sham stimulation (significant interaction between Condition and Time, F_28_ = 3.528, p = 0.043; Fig 3A). Post hoc t-tests revealed a significant increase in mixed percept duration with active cTBS from pre to 5min post (t_14_ = −3.065, p = 0.008) and from pre to 30min post (t_14_ = −2.306, p = 0.037; Fig 3A). There were no effects of cTBS on ocular dominance index or alternation rate (Fig 3B & 3C).

**Fig 3:**
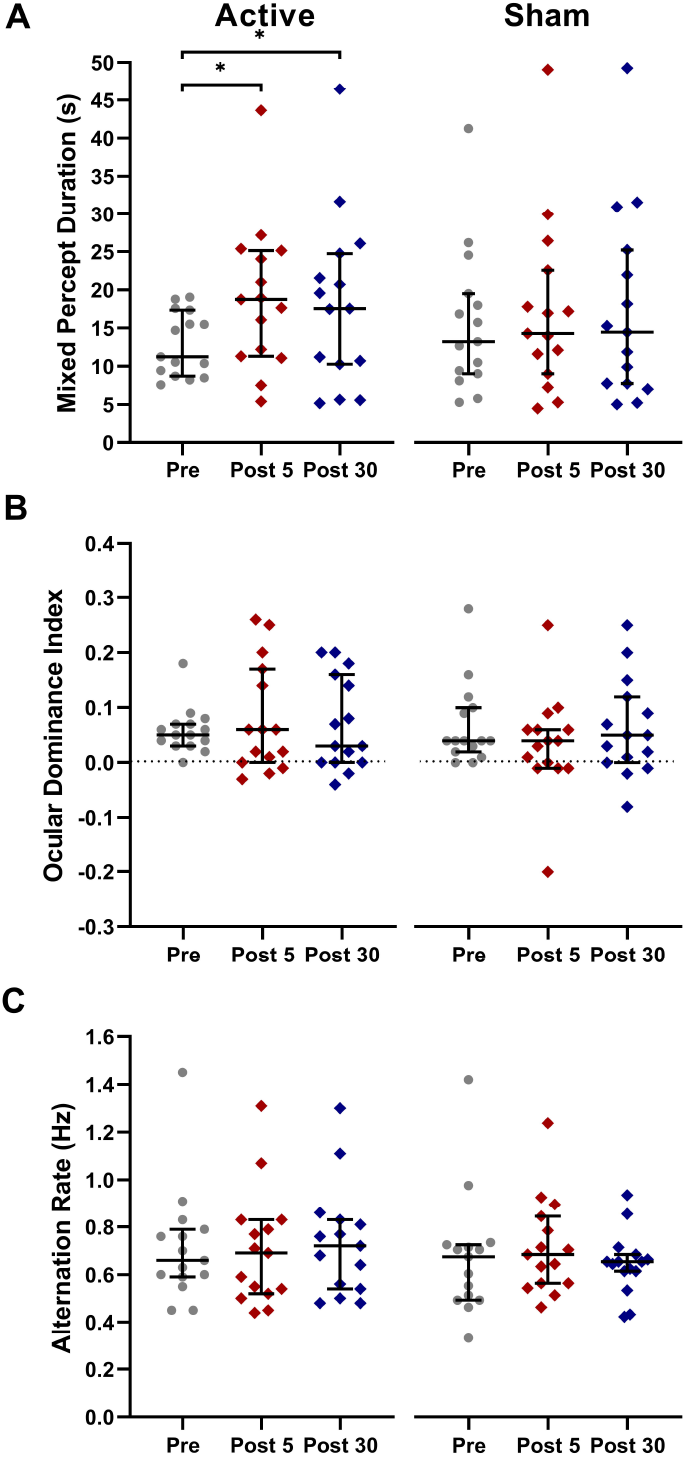
Average time spent in mixed percept (A), ocular dominance indices (B), and alternation rates (C) for 15 participants pre-stimulation, 5 minutes and 30 minutes post cTBS. Error bars = SEM. * p<0.05.

## Discussion

We tested the hypothesis that visual cortex a-tDCS acts to reduce GABA-mediated inhibition within visual cortex as evidenced by reduced binocular rivalry mixed percept duration (29). Mixed percept duration during binocular rivalry has been causally linked to visual cortex GABA concentration through pharmacological antagonism of GABA_a_ and GABA_b_ receptors (29). We also hypothesized that visual cortex cTBS would have the opposite effect to a-tDCS and increase mixed percept duration. This is because while motor cortex a-tDCS has been observed to reduce regional GABA concentration (13,17), cTBS applied to the motor cortex (28) or visual cortex (25) increases GABA concentration.

Contrary to our hypotheses, visual cortex a-tDCS had no effect on mixed percept duration and visual cortex cTBS reduced mixed percept duration, a result that was opposite to the anticipated effect. As expected, neither form of non-invasive brain stimulation altered ocular dominance index or alternation rate during binocular rivalry.

### No effect of a-tDCS on mixed percept duration

The most obvious explanation for the lack of an a-tDCS effect on mixed percept duration is that a-tDCS simply had no effect on the visual cortex at all. Although we certainly can’t rule out this possibility, the vast majority of published studies using the same or similar stimulation parameters over visual cortex have reported a-tDCS effects, including effects that are consistent with reduced GABA-mediated inhibition such as reduced surround suppression (22), reduced lateral inhibition (23) and an equalization of the cortical response to each eye in adults with amblyopia (3,4). Therefore, we also propose a number of alternative explanations.

One explanation is that a-tDCS does reduce visual cortex GABA concentration, but that the primary visual cortex is not the appropriate region to target because a broad network of brain areas that includes the LGN (42), V1 (43) and the prefrontal cortex (44,45) is involved in binocular rivalry. However, an association between GABA concentration and binocular rivalry dynamics has been demonstrated exclusively for the primary visual cortex suggesting that this explanation is unlikely. Furthermore, we observed an effect of visual cortex cTBS on mixed percept duration indicating that visual cortex stimulation can influence binocular rivalry dynamics.

An alternative explanation is that a-tDCS does not act to modulate GABA concentration in visual cortex. Although most MRS studies have reported reduced GABA concentration following a-tDCS of non-visual brain areas, usually motor cortex, not all studies have observed this effect (18,46–48). It is possible that a-tDCS modulation of GABA concentration is highly dependent on stimulation parameters (see Grasso et al., 2020 for a comprehensive review) and/or the distribution of male and female participants within the study population (50–52), all of which differ among previous studies and our own. In addition, it is possible that a-tDCS mechanisms differ between brain regions. For example, Dwyer et al. (53) recently reported no change in GABA or GLX concentration following temporal lobe a-tDCS. In addition, a detailed study of repetitive TMS (rTMS) effects over different brain regions using functional connectivity and computational modeling has recently revealed that the effects of a fixed 1 Hz stimulation protocol differ significantly when the stimulation is applied to different brain regions (54). In particular, occipital 1Hz rTMS induced opposite functional connectivity effects when compared to frontal 1 Hz rTMS. We posit that similar effects occur for a-tDCS and that visual cortex a-tDCS may not influence local GABA concentration in the same way as a-tDCS of motor cortex. An MRS study of visual cortex a-tDCS is required to directly address this question.

### Increased mixed percept duration following visual cortex cTBS

MRS measurements made after visual and motor cortex cTBS have indicated increased visual cortex GABA concentration (25,28), an effect that would be expected to reduce mixed percept duration and increase perceptual dominance (29,55). A possible explanation for our observation of reduced mixed percept duration following visual cortex cTBS relates to changes in the signal-to-noise ratio within visual cortex neural activity (25,26,56). A recent study found that adding noise to the primary visual cortex using transcranial random noise stimulation (tRNS) resulted in a significant reduction in mixed percept duration (56). In other words, an increase of neural noise within the visual cortex increased interocular suppression and therefore reduced mixed percept duration. Similarly, a cTBS-induced reduction in neural noise was proposed by Allen et al. (2014) to explain their observation that visual cortex cTBS improved visual task performance even though the same cTBS protocol increased both phosphene thresholds and visual cortex GABA concentration. Reduced neural noise may also have contributed to the improved visual acuity in adult amblyopic eyes following visual cortex cTBS reported by Clavagnier and colleagues (24).

Of direct relevance to our results, Allen et al (25) suggested that effect of increased cortical inhibition on visual cortex signal-to-noise ratio is dependent on the extent to which inhibition is increased. Applying this idea to binocular rivalry mixed percept duration, a small increase in visual cortex inhibition such as that induced by cTBS may act to reduce neural noise and therefore increase mixed percept duration (56). However, a very large increase in neural inhibition that directly affects excitatory signalling such as that induced by the systemic administration of GABA receptor agonists may reduce mixed percept duration and increase perceptual dominance as reported by Mentch and colleagues (29). In this context, our cTBS results are consistent with generating a small increase in visual cortex inhibition that was sufficient to reduce neural noise but not to substantively alter excitatory signalling. Continuing this line of reasoning, a possible explanation for the null effect of a-tDCS is that a-tDCS increased excitation to a level where the effect of increased neural noise (reduced mixed percept duration) cancelled out the effect of reduced GABA concentration (increased mixed percept duration). This speculation requires objective confirmation of reduced visual cortex GABA concentration following visual cortex a-tDCS.

### Measures of binocular rivalry dynamics

There are differences in binocular rivalry dynamics calculations across previous studies, and subtle differences in definitions. For instance, some studies calculate the proportion of perceptual dominance to mixed percept perception (referred to as perceptual suppression) (29,36), while mean dominance durations calculated as the average duration that a dominant percept lasts in seconds is used by others (34). Our measures were designed to capture any changes in dominance and mixed percepts.

We also measured alternation rates because previous studies have suggested that visual cortex GABA concentration is correlated with alternation rate in young adults (34,36). We did not observe an association between alternation rate and visual cortex a-tDCS or cTBS. Notably, GABA agonists do increase perceptual suppression (i.e. reduce mixed percept duration), but do not consistently influence alternation rate suggesting that different mechanisms may gate alternation rate (29,34).

Overall, our results suggest that the effects of a-tDCS on GABA concentration may differ between the visual cortex and the motor cortex. Further investigation of this question using techniques such as MRS is required to elucidate the mechanisms of visual cortex a-tDCS and help guide the continued development of visual rehabilitation strategies that involve a-tDCS or other forms on non-invasive brain stimulation.

## Acknowledgements

This study was supported by Natural Sciences and Engineering Research Council (NSERC) Grants RPIN-05394 and RGPAS-477166 as well as the Canadian Foundation for Innovation (CFI) Grant 34095.

The funders had no role in study design, data collection and analysis, decision to publish, or preparation of the manuscript.

